# Vertical Variation of the Caterpillar Community in Oak (*Quercus robur*) Canopies

**DOI:** 10.64898/2026.04.07.717053

**Authors:** Lucy M. Morley, Ella F. Cole, Sam J. Crofts, Ben C. Sheldon

## Abstract

**Background:** Understanding how caterpillar communities vary within tree canopies is key to interpreting forest trophic dynamics and responses to environmental change, yet such variation remains poorly quantified due to the challenges of sampling in three dimensions.

**Aims:** We quantified within-canopy heterogeneity in caterpillar densities, diversity, and herbivory and explored relationships with host tree phenology and commonly used ground-based monitoring approaches.

**Methods:** Using direct canopy access, we sampled branches from lower, middle, and upper canopy strata of 34 mature pedunculate oaks (*Quercus robur*) in Wytham Woods, UK, during the spring abundance peak over three consecutive years (2023-2025). We tested for vertical stratification in caterpillar community metrics, examined patterns in early instar distributions at emergence, assessed associations with host tree phenology across spatiotemporal scales, and evaluated how well ground-based methods (water and frass traps) reflect canopy communities.

**Results:** Vertical stratification was modest but varied among years: densities and species richness increased with canopy height in 2023, decreased in 2024, and were uniformly low across strata in 2025. Although within-crown budburst timing varied systematically, with upper branches bursting approximately two days earlier than lower branches, tree phenology did not explain within- or between-year variation in caterpillar communities. Frass trap data correlated moderately well with canopy caterpillar densities, whereas water traps showed weaker and less consistent relationships, reflecting behavioural and methodological biases.

**Conclusions:** Caterpillar communities showed no consistent patterns of vertical stratification across years, instead they are shaped more strongly by inter-annual and tree-level variation. Integrating targeted canopy sampling with scalable ground-based proxies could greatly improve monitoring of arboreal Lepidoptera and inform studies of trophic synchrony and woodland resilience under environmental change.

## Introduction

Tree canopies are structurally complex habitats that support diverse arboreal insect communities. In temperate woodlands, vertical gradients in microclimate (e.g. light intensity, temperature, humidity), leaf structural morphology and chemistry, predation risk, and competition all contribute to generating fine-scale spatial heterogeneity within individual tree crowns. This creates diverse ecological niches that can promote species coexistence, specialisation, and high bio-diversity (Seifert et al., 2020; Xing et al., 2023). In pedunculate oak (*Quercus robur*), this heterogeneous canopy supports a particularly abundant and diverse community of phytophagous Lepidopteran larvae (hereafter “caterpillars”), with over 250 species recorded on *Q. robur* in the UK (Mitchell et al., 2019). As primary consumers, these caterpillars form a key trophic link between trees (producers) and secondary consumers (such as insectivorous passerine birds) in the well-studied tri-trophic woodland system. Much of this work explores the causes and consequences of changing seasonal timing (phenology) in trophic interactions, often relying on estimates of caterpillar abundance and species composition derived from monitoring methods that are ground-based or restricted to accessible lower branches (van Asch & Visser, 2007; Charmantier et al., 2008; Shutt et al., 2019; Burgess et al., 2018; Macphie et al., 2025). However, the extent to which these approaches accurately describe the diversity and phenology of the community in trees, or capture within-canopy variation in caterpillar community dynamics, remains poorly understood, largely due to the logistical challenges of sampling across vertical strata.

Among oak-feeding Lepidoptera, the winter moth (*Operophtera brumata*, Geometridae) and green oak tortrix (*Tortrix viridana*, Tortricidae) will be explicitly addressed in this study because they are abundant species within the spring caterpillar community at our study site. Both species exhibit cyclical population dynamics, interspecific competition, and can cause extensive defoliation through herbivory (Hunter & Willmer, 1989; Hunter, 1990; Hunter, 1992). Winter moth in particular are exemplified in the simplified tri-trophic oak – winter moth – great tit (*Parus major*) food chain, where their abundances are typically measured by water traps on the ground (Hinks et al., 2015; Charmantier et al., 2008). Phenological synchrony between oak budburst and larval emergence time is thought to be critical for early instar winter moth survival, but climate-driven mismatches can reduce caterpillar fitness and abundance (van Dongen et al., 1997; van Dis et al., 2023). As a key food resource for insectivorous birds, reduced winter moth caterpillar availability may subsequently affect reproductive fitness of secondary consumers, subject to their own phenotypic plasticity (Visser & Holleman, 2001; Visser et al., 2006; Charmantier et al., 2008). Understanding how the vertical distribution of caterpillar species varies within individual tree canopies, as well as between trees, in relation to host tree phenology would therefore be insightful for both assessing previous estimates of caterpillar communities, as well as predicting trophic dynamics under environmental change.

### Drivers of vertical stratification

Vertical stratification in caterpillar communities may arise from species-specific life history traits. In winter moths, flightless adult females ascend tree trunks during winter and oviposit progressively with height, preferring to lay on the uppermost branches (Visser & Holleman, 2001; Watt et al., 1992; Graf et al., 1995). In conjunction with fine-scale temperature variation affecting egg development rate, this could generate gradients in egg density and hatching phenology (Hibbard & Elkinton, 2015; van Dis et al., 2021). In contrast, species with fully flighted females, such as *T. viridana*, may be less likely to exhibit any oviposition-driven stratification. Such interspecific behavioural differences provide a mechanistic basis for expecting vertical heterogeneity, particularly of winter moth caterpillars, in our study system.

Once hatched, spatiotemporal variation in leaf traits within tree crowns may also influence caterpillar distributions. Upper canopy (i.e. more sun-exposed) leaves typically differ from shaded lower leaves in thickness, toughness, and chemical composition (water, nitrogen, chlorophyll and defensive compounds; Yamasaki & Kikuzawa, 2003; Volf et al., 2022; Gardner et al., 2022; Hakimara & Despland, 2025a; Blair, 2024). Such variation can affect leaf nutritional quality and palatability as they develop, although reported effects on subsequent caterpillar performance are mixed across systems and between or within tree scales (Hemming & Lindroth, 1995; Eisenring et al., 2021; Hakimara & Despland, 2025b; Stiegel et al., 2017; Ruhnke et al., 2009). For example, autumnal moth caterpillars (*Epirrita autumnata*, Geometridae) performed better when fed on leaves taken from upper branches of mountain birch trees (*Betula pubescens* ssp. *tortuosa*) compared to lower leaves (Suomela et al., 1995a). Performance also varied among leaves from different compass directions and hierarchical levels (ramets, branches, shoots) within trees, not just vertically and were followed up with leaf biochemical analysis (Suomela & Nilson, 1994; Suomela et al., 1995b). These findings indicate there are effects of fine-scale resource quality, mediated by leaf traits, on caterpillar fitness that could determine their within-crown distributions.

Leaf traits are also closely linked to phenology as leaves mature over the growing season, so within-crown variation in budburst phenology could reinforce vertical caterpillar stratification. Empirical work by Feeny (1968) showed that decreasing nitrogen, and increasing defensive tannins and leaf toughness, make older *Q. robur* leaves less palatable, corresponding to lower winter moth caterpillar densities. Other studies suggest that younger nitrogen-rich leaves generally have better caterpillar performance (Stamp & Bowers, 1990; Coley et al., 2006; Molleman et al., 2022). In *Quercus alba*, canopy branches (>3m high) came into leaf six days earlier than understorey (<3m high) branches, thereby creating spatiotemporal variation in food resource availability (Barber & Fahey, 2015). Caterpillars can respond through selective foraging or dispersal, both within and between trees, to track optimal resources, e.g. *Zeiraphera canadensis* (Tortricidae) larvae disperse upwards to track acropetal bud development in white spruces (*Picea glauca*; Carroll & Quiring, 1994). The extent to which within-canopy phenological variation structures whole caterpillar communities, particularly in *Q. robur*, remains underexplored.

### Evidence for vertical variation in canopy caterpillar communities

Vertical stratification of arboreal Lepidoptera has been reported from temperate and tropical forests, although patterns are inconsistent among systems and studies have rarely been conducted across multiple years. Some studies focus on single species, e.g. forest tent caterpillars (*Malacosoma disstria*, Lasiocampidae) on quaking aspen (*Populus tremuloides*; Batzer et al., 1995), while other studies describe community-level variation. Seifert et al. (2020) collected 3,892 caterpillars from 215 species on felled deciduous trees, finding that species composition, but not caterpillar density, differed among understorey, midstorey, and canopy strata. Despite reduced richness and overall diversity, shelter-builders and increased specialisation were favoured toward the upper canopy (Seifert et al., 2020). In the same system, Blair (2024) observed peak caterpillar density in the mid-canopy, correlated with leaf thickness and feeding guild, with reduced upper canopy herbivory likely due to lower leaf nutritional quality. Similar guild-specific vertical patterns of greater mid-canopy caterpillar densities were also reported in a European forest (Šigut et al., 2018). In US *Q. alba* and *Q. velutina*, Corff and Marquis (2001) found a 60% increase in insect herbivore density in the canopy (15-20m high) than understorey (<2.5m) strata, but only at one of six sampled timepoints. However, species richness was always higher by 5-20% in the understorey with differing compositions between strata, partially mediated by leaf nutritional traits (Corff & Marquis, 2001).

Similar guild-specific or structural effects have been observed in tropical forests, where stratification patterns vary by taxa and location (Schulze et al., 2001; Brehm, 2007; Neves et al., 2013; Ashton et al., 2015; Finnie et al., 2024). Collectively, these studies reveal substantial vertical variation in caterpillar abundance, diversity, and composition, but the direction and magnitude of effects are context dependent. Few studies address fine-scale within-canopy strata rather than canopy versus understorey, extend across multiple years to examine temporal consistency, or consider phenological dynamics.

### Implications for ground-based monitoring

Common approaches for monitoring arboreal caterpillars, such as frass traps, water traps, and branch beating, are ground-based or restricted to accessible lower branches (Visser et al., 2006; Hinks et al., 2015; Shutt et al., 2019). These methods collapse three-dimensional canopy complexity into two-dimensional samples, which may not accurately reflect canopy communities so raises questions about their representativeness. Water traps are likely to be biased towards species that descend to the ground to pupate in the soil rather than those pupating within canopies. Frass traps estimate community biomass by collecting faecal pellets but lack species-level resolution (Rytkönen et al., 2018). Meanwhile, branch beating actively collects live individuals rather than a passive accumulation between sampling intervals but may be difficult in trees lacking accessible branches, or underestimate the abundance of leaf-rolling species. Tests by Zandt (1994) on *Q. petraea* and *Q. robur* suggested that temporal (i.e. whether comparing seasonal totals or individual sampling dates) and weather-related variability can reduce the agreement of these methods. More destructive techniques (e.g. whole tree felling: Seifert et al., 2020; Finnie et al., 2024; and insecticide fogging: Summerville et al., 2003; Müller et al., 2017) are unsuitable for phenological monitoring across time. Such limitations mean we cannot therefore assume that ground-based field methods accurately describe within-canopy caterpillar activity. Directly accessing the canopy using mobile elevated work platforms (MEWPs) offers a practical alternative for resolving vertical sampling of live caterpillar communities to enhance our understanding of community dynamics (Šigut et al., 2018).

Here we conducted a field study in a temperate deciduous woodland to quantify fine-scale vertical variation in caterpillar communities hosted by oak trees and understand potential phenological drivers. We carried out MEWP-based branch sampling of 34 mature *Q. robur* trees in Wytham Woods over three consecutive springs (2023-2025), complemented with intra-annual and ground-based sampling, to test how vertical structure may shape caterpillar community dynamics and the reliability of common monitoring approaches.

Specifically, we aimed to:

1. Quantify variation in caterpillar density, diversity, and herbivory across lower, middle, and upper canopy strata.
2. Test whether this variation is linked to host tree phenology, measured using bud scoring and remotely sensed NDVI.
3. Evaluate how well ground-based methods (water and frass traps) reflect within-canopy caterpillar community measures.

## Methods

### Study site and tree selection

This study was conducted in Wytham Woods, Oxfordshire, UK (51°46’N, 1°20’W, National Grid Reference SP4608), a long-term ecological research site dominated by temperate deciduous woodland. We selected 34 mature pedunculate oak (*Quercus robur*) trees along an arterial path that were accessible using a mobile elevated work platform (MEWP). Trees were chosen opportunistically based on accessibility, safety of access, and spatial spread across the woodland (see map in **SI Figure 1**).

### Branch sampling

At each tree, we sampled three vertical canopy strata, defined as the relative lower, middle and upper thirds of the total crown height. At each level, a terminal branch approximately 50cm in length was enclosed in a large plastic bag and then cut from the tree using secateurs. In 2023, we sampled two branches from 24 canopy levels across 11 trees to test within-strata repeatability and randomly retained one branch from the pair for the main analyses. We measured branch height above ground (to the nearest metre) using a tape measure.

Primary branch sampling occurred during the spring peak of arboreal caterpillar abundance, over one week each year: 15^th^ to 21^st^ May 2023 (n=96 branches), 13^th^ to 18^th^ May 2024 (n=98 branches), and 12^th^ to 17^th^ May 2025 (n=91 branches). Sampling weeks were aligned to the same calendar week across years, although phenological timing shifted slightly among years so this may not reflect identical points of the caterpillar phenological distribution (see Discussion). Sampled branch heights ranged from 2-20m. One tree was not sampled in 2025 due to obstruction by a fallen tree.

### Laboratory processing of branches

On the same day, we shook the sealed bags to dislodge invertebrates before removing the branch to extract all caterpillars. We simultaneously stripped all the leaves to ensure no caterpillars remained undetected and weighed fresh leaf mass to the nearest gram. All caterpillars were identified where possible using field guides (Henwood et al., 2020; Porter, 2010) and suggestions from the ObsIdentify app (Observation International, 2025). Samples were then preserved individually at -20°C.

Across years, 92.63% of larvae were identified to species or morphospecies level of Lepidoptera. An additional 1.72% were classified as Symphyta (sawfly larvae), and 5.65% remained unidentified, due to having no clear defining characteristics (e.g. early instars or damaged), which were included in total counts but excluded from species richness and diversity metrics (a comparable 5.5% of individuals were unidentified by Seifert et al., 2020). We calculated densities of total caterpillars, winter moths (*Operophtera brumata*), green oak tortrix larvae (*Tortrix viridana*), and species richness, all standardised per 100g leaf mass to account for variation in branch size (leaf mass range = 15-265g; by year: 2023 = 30-160g, 2024 = 15-105g, 2025 = 35-265g). We also calculated Shannon diversity index values per branch as a scale-independent measure of diversity accounting for species number and abundance.

We evaluated the assumption that species richness scales linearly with sampling effort by plotting raw species richness against branch leaf mass (**SI Figure 2**). Species richness increased weakly but approximately linearly with leaf mass in all years, with no evidence of saturation across the range of sampled leaf mass, supporting the use of species richness density as counts standardised by leaf mass rather than rarefaction-based approaches.

To estimate herbivory, we randomly selected 10 leaves per branch. In 2023, percentage area consumed was visually categorised into 5% intervals (0-5%, 6-10%, 11-15%, 16-20%, 21-25%, and 26-30%) then we took the midpoint of each category multiple by the number of leaves and divided by 10 to calculate a single mean value. In 2024 and 2025, herbivory was quantified more precisely using the LeafByte mobile app, which measures percentage of total leaf area consumed from scale-referenced images (Getman-Pickering et al., 2020). From both of these methods, we used mean percentage leaf area consumed as our measure of herbivory. Individual leaf mass to the nearest 0.01g was also recorded.

### Intra-annual branch sampling in 2024

To assess within-year dynamics, additional sampling of the same trees occurred in 2024 between 10^th^ to 16^th^ April (n=92 branches). This was to coincide with oak budburst to quantify densities of newly emerged early instar caterpillars, scaled to 100g bud mass, but these were not visually identifiable to species. We also scored the branch phenology, as it varied between trees and might influence caterpillar presence, using a bud development scale from 1 to 7 (1 = little to no bud swelling, 4 = budburst, 7 = fully expanded leaf) established in Hinks et al. (2015; further used in Cole & Sheldon (2017) and Morley et al. (2025)).

### Host tree phenology via budburst scoring and remotely sensed NDVI

To explore the influence of within-crown phenological variation on caterpillar communities, in 2024 we monitored bud progression of focal trees in the field using the key described above. We scored 12 sections of each tree (four horizontal quarters of the lower, middle and upper thirds) every 3-4 days (from 26^th^ March to 23^rd^ May) then calculated the whole crown mean score across the 12 sections and the mean of each canopy level from its four quarters. We standardised scores to between 0 (stage 1) and 1 (stage 7) and fitted logistic curves, controlling for observer effects, to extract the date on which the curve crossed y=0.5 (stage 4) as the budburst date for each whole tree crown or stratified canopy level (see Morley et al. (2025) for further details).

Budburst data was only available for 2024, so to relate caterpillar metrics to phenological variation of host tree crowns across years, we collected multispectral drone imagery using WingtraOne Gen II drones equipped with RedEdge-P cameras in all three years. Flights were conducted every 3-4 days between 26^th^ March and 9^th^ June 2023, 28^th^ March and 14^th^ June 2024, and 24^th^ March and 4^th^ June 2025. We stitched the orthomosaics, with PPK GPS corrections, for each flight in Pix4Dfields then extracted NDVI values for the core 75% of each individual manually delineated tree crown across the time series. One of the 34 tree crowns could not be delineated due to overlap of neighbouring crowns. NDVI values were standardised between each tree’s seasonal minimum and maximum then calculated the half-NDVI date (date of 50% of the seasonal maximum) as an inter-annual measure of whole crown phenology. For detailed image acquisition and processing protocols, and validation of the NDVI-based phenology metric as an accurate proxy for oak budburst, see Morley et al. (2025).

### Ground-based measures of caterpillar communities

To relate canopy-level measures to ground observations, we placed frass traps and water traps beneath each focal tree in 2024 and 2025. Frass traps provided an estimate of caterpillar community biomass, while water traps gave abundance and diversity metrics. Ground sampling was not available for 2023 due to MEWP-sampled trees being selected mid-season according to logistical constraints of the machinery, whereas water traps need to be put out in early spring.

Frass traps consisted of a muslin cloth attached to a 50cm x 50 cm (0.25m^2^) wire frame one metre above the ground on bamboo cane supports, with a stone in the centre to accumulate matter on the cloth. Traps were set up three days prior to branch sampling, then contents were scraped into envelopes, oven-dried at 50°C for 24 hours to remove moisture content and sieved to remove leaf litter so frass could be weighed to the nearest 0.01g.

We also placed water traps beneath focal trees to intercept caterpillars falling from the canopy. These consisted of 100cm x 55cm (0.55m^2^) plastic trays filled with water to a depth of approximately 10cm which we dredged every three days from 28^th^ April to 3^rd^ June 2024, and 2^nd^ May to 2^nd^ June 2025 (the peak duration was shorter in 2025). Collected caterpillars were counted and identified to species level (where possible). Sampling dates were part of wider spatial sampling across the site, so caterpillar counts may have been on the same day as branch sampling or ±1 day.

### Statistical analyses

All statistical analyses were conducted in R v4.5.1 (R Core Team, 2025). We used Bayesian generalised linear mixed models (GLMMs; brms package; Bürkner, 2017) to test the effects of canopy level, sampling year, and tree phenology on six metrics of caterpillar communities: total caterpillar density, density of winter moth caterpillars, density of green oak tortrix caterpillars, species richness density, Shannon diversity index (alpha diversity; density-independent measure of biodiversity accounting for both abundance and species number), and mean percentage leaf area consumed as herbivory. Densities were raw count data scaled per 100g of fresh leaf mass to account for variation in branch size. All density and Shannon index responses were zero-inflated, so were modelled using hurdle lognormal models to account for zero values (probability of caterpillar absence) and non-zero densities (conditional on presence). Herbivory, which contained no zero values, was modelled using a lognormal distribution. All models included the random effect of tree identity to account for repeated sampling of branches within trees and across years. We checked convergence and model fit with posterior predictive checks, and effects were considered significant if 95% credible intervals excluded zero. Coefficients in the results are presented on the output link scale, i.e. not back-transformed, for simpler interpretation (i.e. positive coefficients are increasing effects, negative coefficients are decreasing effects, rather than back-transformed to multiplicative effects).

We also quantified repeatability using intraclass correlation coefficients (ICC) across the six response variables, from the trees which we had sampled two branches from the same stratum in 2023 (Gamer et al., 2019). To test how well ground methods reflected caterpillar communities in tree canopies, we calculated Spearman’s rank correlations (r_s_) between branch and ground-based measures (frass and water traps). Frass data was mass accumulated for three days prior to branch sampling, whereas we tested water trap counts corresponding to the nearest collection date of a wider sampling schedule (the same or ±1 day) as well as the seasonal total.

## Results

### Caterpillar counts and species compositions

Across the three sampling years, a total of 1,221 caterpillars were collected from the 34 oak trees during the main May sampling periods, representing 41 species (32 identified to species level and 9 morphospecies) from six Lepidopteran family groups, as well as a number of sawfly larvae (order Symphyta). The number of larvae collected differed between years: 512 individuals in 2023, 505 in 2024, and a marked reduction to 204 in 2025. When standardised by sampling effort (total larvae per 100g leaf mass), overall densities were highest in 2024 (10.73 larvae), intermediate in 2023 (6.31 larvae), and substantially lower in 2025 (2.45 larvae), indicating that the reduced abundance in 2025 reflects a genuine decline in caterpillar density rather than differences in the number of branches or leaf mass sampled (see Methods).

Species composition varied among canopy levels and years (**Figure 1**). In all years, winter moth (*Operophtera brumata*, Geometridae) was consistently the most abundant species, though its relative abundance declined slightly in 2024 and 2025. Green oak tortrix (*Tortrix viridana*, Tortricidae) was the second most abundant species in 2023 and 2024, but only a single larva was recorded in 2025, when other Geometridae became more common. These two dominant species, highlighted separately from their family groups in **Figure 1**, together accounted for approximately half of all larvae in 2023 and 2024, and winter moth for one-third of larvae in 2025. Other frequent taxa included Noctuidae, other Geometridae and other Tortricidae, with low frequencies of other Lepidopteran taxa (Depressariidae, Eredibae, and Lycaenidae) and a small number of Symphyta larvae. A total of 69 individuals (5.65%) could not be visually identified to family level. Overall, community composition showed only minor variability across years and canopy levels, predominantly reflecting shifts in the relative abundance of winter moth and green oak tortrix larvae.

**Figure 1.**
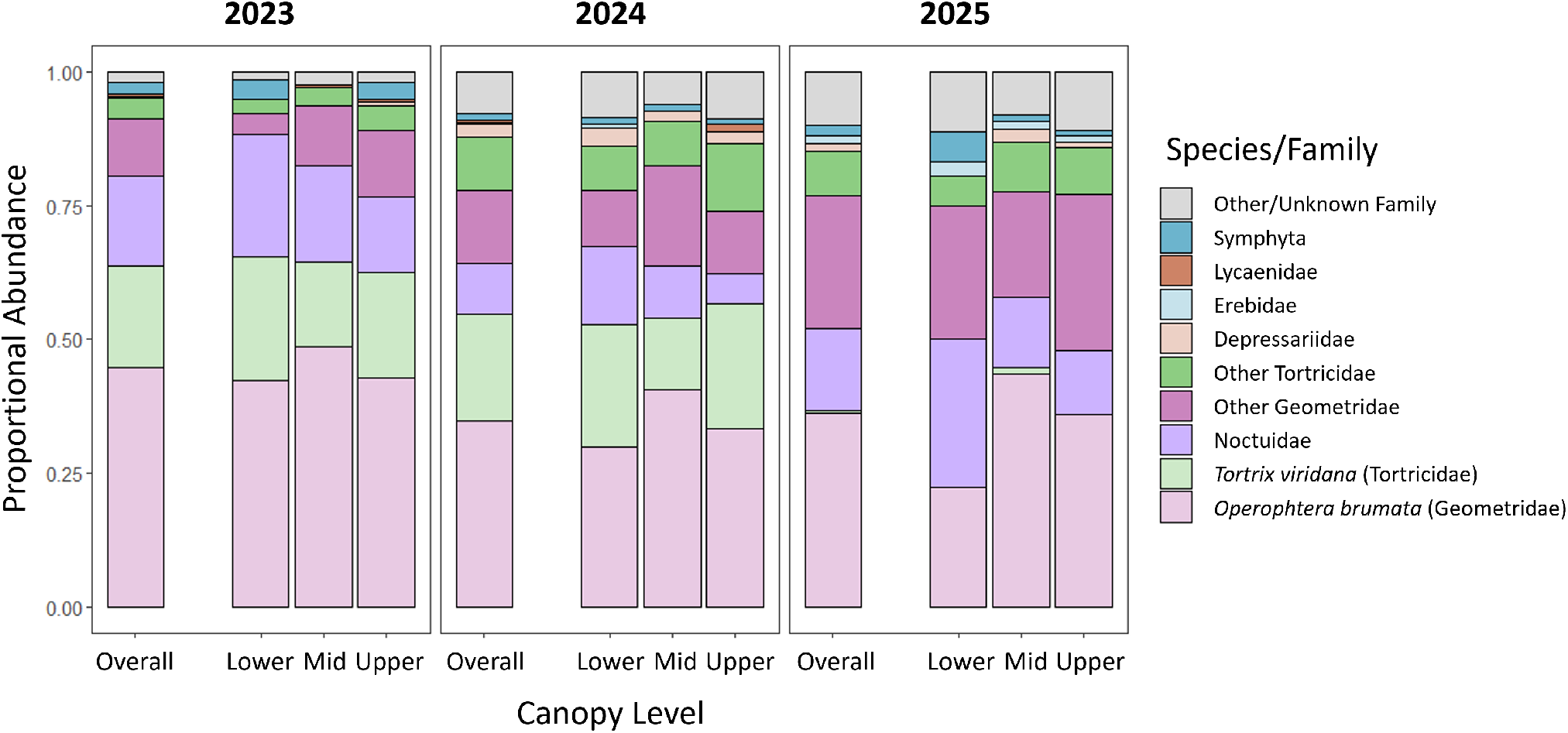
Proportional abundance of caterpillar families and two key species recorded during May branch sampling of 34 oak trees in 2023–2025, shown for the overall total across branches and by canopy level branches came from.

### Repeatability within canopy levels

To assess repeatability of caterpillar community metrics within canopy levels, a subset of trees in 2023 had two branches sampled per some canopy levels (n=24 pairs of branches from 11 trees). Total caterpillar density, winter moth density, species richness density, and herbivory (mean percentage leaf area consumed) all showed moderate repeatability (ICC, R=0.46–0.52, all p<0.01). Shannon diversity index showed lower, but only marginally non-significant, repeatability (R=0.33, p=0.052), while green oak tortrix density was not repeatable within canopy levels (R=0.12, p=0.265). These results indicate moderate consistency in most caterpillar community aspects between branches within the same canopy level for this subset of trees.

### Effect of canopy level across years

Total caterpillar density varied significantly among canopy levels and years (**Figure 2a**). In 2023, mean density increased with height, with the upper canopy (8.50 caterpillars per 100g leaf mass) having significantly higher densities than both lower (4.27) and mid (5.95) branches (post-hoc emmeans contrasts: upper–lower +4.20, 95%CI [1.67, 6.96]; upper–mid +2.54, 95%CI [0.03, 5.32]; both p<0.05). In contrast, 2024 showed the reverse pattern, with densities highest in the lower canopy (12.97) and significantly reduced in the upper canopy (7.76; upper–lower -5.18, 95%CI [-9.86, -0.95], p<0.05) with intermediate mid-canopy density (10.96). No significant differences among levels were detected in 2025, when densities were uniformly low (lower=2.18, mid=2.47, upper=2.34).

**Figure 2.**
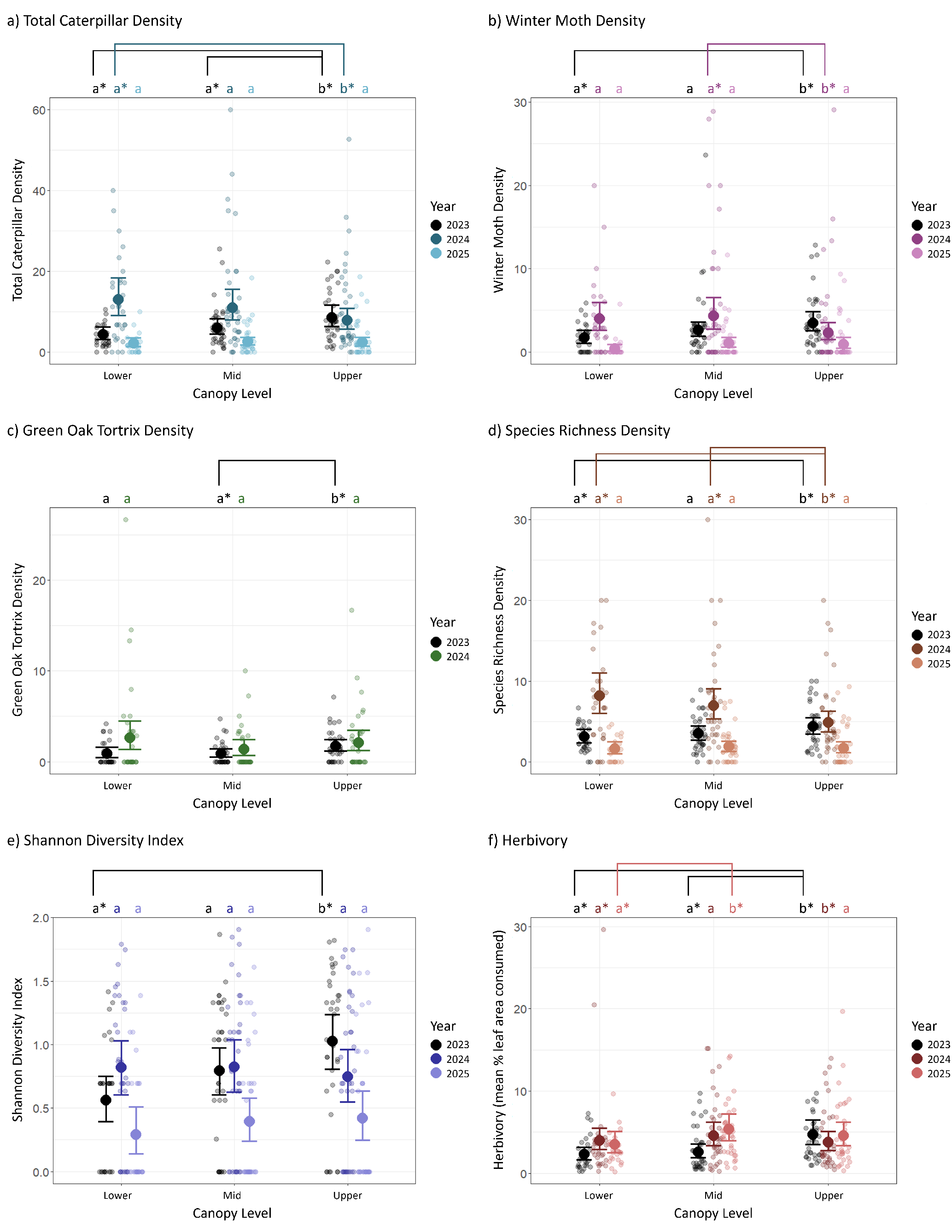
Conditional effects plots of the modelled interaction between canopy level (lower, mid, upper third) and year on caterpillar community metrics across n=285 branches from 34 oak trees, sampled from Wytham Woods, UK, in 2023-2025. Large points and error bars show means and 95% credible intervals of combined hurdle lognormal model estimates, averaged across the random effect of tree identity. Density measures (a-d) are abundance counts scaled to 100g leaf mass, whereas (e-f) are scale-independent. In (c), 2025 was excluded from modelling due to only one individual green oak tortrix larva being recorded. Letters and asterisks denote significant (p<0.05) within-year pairwise differences among canopy levels based on model estimated marginal means.

These patterns reflected significant effects on both model components: the probability of caterpillar absence (zero hurdle) and densities when present (non-zero part; see **SI Table 1** for full model outputs). Conditional on presence, densities were significantly higher in the upper canopy than lower canopy in 2023 (β=0.61, 95%CI [0.21, 0.98]), and higher on low branches in 2024 than 2023 (β=1.14, 95%CI [0.75, 1.54]). However, this was offset by a negative interaction reducing upper canopy densities in 2024 (upper x 2024: β=- 1.17, 95%CI [-1.70, -0.64]). The zero component indicated a lower probability of caterpillar absence in the upper canopy (β=-9.86, 95%CI [-30.65, -0.42]) relative to a 6.1% baseline in 2023 low branches, but a higher probability of absence on them in 2025 (β=2.15, 95%CI [0.55, 4.14]), particularly more so in the upper canopy that year (upper x 2025: β=9.89, 95%CI [0.37, 31.15]). Collectively, these results indicate that total caterpillar density shifted both vertically and temporally, with strong upper canopy dominance in 2023, reversed patterns in 2024, to uniformly low densities in 2025.

Winter moth densities showed similar temporal and vertical patterns to the overall community (**Figure 2b**), being higher in the upper canopy than lower canopy in 2023 (upper–lower +1.80, 95%CI [0.59, 3.04], p<0.05), but lower than mid-level branches in 2024 (upper–mid -1.98, 95%CI [-4.12, -0.11], p<0.05). Again there were no differences among levels in the low abundance year of 2025. Conditional effects showed higher densities on low branches in 2024 (β=0.73, 95%CI [0.29, 1.15]), counteracted again by a significant negative upper canopy interaction (upper x 2024: β=-0.75, 95%CI [-1.31, -0.16]). Probabilities of absence were lower in mid (β=-1.53, 95%CI [-2.92, -0.23]) and upper (β=-1.91, 95%CI [-3.50, -0.52]) canopy levels in 2023, but increased in 2024 (mid x 2024: β=2.10, 95%CI [0.44, 3.92]; upper x 2024: β=2.47, 95%CI [0.70, 4.39]), and again in 2025 (β=2.06, 95%CI [0.85, 3.40]), all relative to low branches in 2023, consistent with the reduced densities shown in **Figure 2b**.

Green oak tortrix larvae also showed generally higher densities in 2024 than 2023 on low branches where present (β=0.98, 95%CI [0.41, 1.53]), although abundance dropped sharply in 2025 (only one larva was recorded, from a mid-level branch) so that year was excluded to enable model convergence. The only significant zero component effect was a lower probability of absence in the upper canopy in 2023 (β=- 1.50, 95%CI [-2.61, -0.44]) compared to low branches, consistent with the higher mean density in **Figure 2c** (2023 upper–mid +0.85, 95%CI [0.08, 1.56], p<0.05), but there were no level differences detected in 2024.

Caterpillar species richness density, i.e. number of caterpillar species per 100g leaf mass, also varied among canopy levels and years (**Figure 2d**). Richness increased with canopy level in 2023 (upper–lower +1.25, 95%CI [0.02, 2.43], p<0.05), but declined in 2024 (upper–lower -3.29, 95%CI [-6.05, - 0.88]; upper–mid -2.08, 95%CI [-4.02, -0.07]; both p<0.05), with no level differences in 2025. Model effects showed that conditional on presence, richness was higher in 2024 than 2023 on low branches (β=1.05, 95%CI [0.72, 1.37]) but reduced in the upper canopy that year (upper x 2024: β=-0.84, 95%CI [-1.27, -0.39]). As with total caterpillar density, the upper canopy had a lower probability of species absence (β=- 9.97, 95%CI [-31.48, -0.29]), relative to the 5.8% baseline in 2023 low branches, but absence probabilities increased in 2025 (β=2.21, 95%CI [0.54, 4.28]), especially in the upper canopy (β=10.12, 95%CI [0.27, 31.50]).

Shannon diversity index (**Figure 2e**) also increased with canopy level in 2023 (upper–lower +0.46, 95%CI [0.19, 0.72], p<0.05) but showed no significant differences in 2024 or 2025. Compared to low branches, the model confirmed again a positive upper canopy effect in 2023 (β=0.34, 95%CI [0.13, 0.56]), and a marginal increase on them in 2024 (β=0.23, 95%CI [0.01, 0.46]) that was offset by a negative upper canopy interaction (β=-0.35, 95%CI [-0.66, -0.05]). Combined, these effects result in the small differences among level means in 2024 being non-significant (**Figure 2e**). The reduced Shannon values in 2025 were explained by a higher probability of zero values (β=1.38, 95%CI [0.26, 2.59]) than on low 2023 branches, meaning a greater prevalence of branches either lacking any caterpillars or containing any number of individuals belonging to only one species.

Finally, herbivory (mean percentage leaf area consumed on 10 leaves per branch; **Figure 2f**) increased with canopy level in 2023 (upper–lower +2.44, 95%CI [1.00, 3.96]; upper–mid +2.13, 95%CI [0.84, 3.70]; both p<0.05) but did not differ among levels in 2024. In 2025 however, herbivory was the only response variable with significant differences among levels, being higher on mid-level branches (mid–lower +1.80, 95%CI [0.04, 3.64], p<0.05). Model effects reflected these patterns, with higher herbivory on upper than lower branches in 2023 (β=0.73, 95%CI [0.35, 1.11]) and higher on them in 2024 (β=0.54, 95%CI [0.15, 0.93]), though this was again moderated by a negative interaction in the upper canopy (upper x 2024: β=-0.77, 95%CI [-1.31, -0.22]). A small but significant increase in herbivory on lower branches existed in 2025 compared to 2023 (β=0.44, 95%CI [0.02, 0.85]) that was not apparent in any other caterpillar response variable.

### Intra-annual caterpillar density

The density of early instar larvae collected from branch samples in April 2024 did not differ significantly among canopy levels (**Figure 3**). Mean early instar densities, scaled per 100g bud mass, were estimated at 10.7 larvae for lower branches, increasing slightly to 15.3 for mid-level and 16.8 for upper branches. Neither the zero hurdle nor non-zero components of the model were significant (see **SI Table 2** for full model output). Tree identity and bud stage were included as random effects and explained a substantial proportion of variation in early instar density, as indicated by the difference between conditional and marginal R^2^ values (36.2-2.3=33.9% attributed to random effects).

### Host tree phenology – whole-crown versus stratified budburst

Host tree phenology may contribute to the substantial variance in caterpillar community metrics explained by tree identity in the previous models. To test this, we monitored bud-to-leaf progression in 2024 and estimated budburst dates for the whole tree crown and b) each canopy level strata. The mean budburst date for the 34 whole crowns was day 101.04 (10^th^ April, SD=2.99). When modelled as a function of canopy level (linear mixed model), the stratified budburst data revealed significant within-tree differences in budburst timing. Lower branches burst on average on day 101.92, whereas mid and upper branches were significantly earlier by 1.03 days (95%CI [-1.67, -0.36]) and 2.18 days (95%CI [-2.82, -1.55]) respectively (**Figure 4**). The difference between mid and upper branches was also significant (upper–mid -1.14, 95%CI [-1.78, -0.61]). Tree identity accounted for the majority of variance in budburst timing, explaining 77.1% of the modelled variation (difference between conditional R^2^=84.8%, and marginal R^2^=7.7%). This indicates strong tree-level differences in phenology, where early-bursting trees tend to be consistently early across all canopy levels (see **Figure 4**).

To assess how host tree phenology influence caterpillar communities within canopies, we fitted pairs of models for each of the six response variables, using either budburst stratified by canopy level or whole-crown budburst as predictors, as well as the canopy level interaction. Model comparisons using leave-one-out information criterion (LOOIC) revealed no substantial improvement in fit for either approach (ΔLOOIC <5 in all cases) and the proportion of variance explained (R^2^) was similar between model pairs (see **SI Table 3** for model comparison statistics). Therefore, stratified budburst did not meaningfully improve model performance.

Model estimated effects were non-significant in eleven of the twelve models. The only exception was the stratified bud-burst model of green oak tortrix density. When caterpillars were present, densities remained lower on mid-level branches than low branches (β=-18.13, 95%CI [-35.12, -0.72]) but showed a borderline increase with later budburst (β=0.17, 95%CI [0.00, 0.34]). In the hurdle component, the probability of absence was higher in upper than lower branches (β=47.63, 95%CI [0.72, 103.72]) but decreased slightly with later budburst (β=-0.48, 95%CI [-1.04, -0.01]). Together these results suggest that within-year variation in budburst timing – whether measured at different canopy level strata or the whole-crown scale – did not strongly influence caterpillar community patterns within oak canopies in 2024.

### Host tree phenology – inter-annual tree phenology via NDVI

We subsequently analysed inter-annual phenological effects using drone-derived half-NDVI date as a proxy for whole-crown budburst. Half-NDVI and whole-crown budburst dates correlated well in 2024 for this subset of trees (r_s_=0.61, p<0.001), therefore NDVI provided a consistent phenology measure in 2023 and 2025 when direct budburst observations were not available.

Tree phenology advanced substantially across years. The mean half-NDVI date of the 33 oaks was 120.2 (30^th^ April; SD=2.60) in 2023, 114.7 (SD=6.10) in 2024, and 104.6 (SD=3.17) in 2025 (one tree was excluded from NDVI analyses as its crown was unable to be delineated). We included year only as an additive effect in NDVI-derived phenology models to control for the inter-annual variation without over-complicating the half-NDVI date by canopy level interaction testing whether phenological effects differed among canopy strata.

However, there was little consistent evidence that NDVI-derived tree phenology influenced within-canopy caterpillar metrics. A borderline positive effect of half-NDVI date on total caterpillar density when present was observed but credible intervals just overlapped zero (β=0.04, 95%CI [-0.00, 0.07]). The only clearer phenological signal occurred for green oak tortrix, where later tree phenology corresponded to a reduced probability of caterpillar absence (i.e. increased likelihood of occurrence; β=-0.14, 95%CI [-0.27, -0.02]). No significant differences among canopy levels persisted in these models, reflecting likely increased uncertainty and complexity of including an additional between-tree variable rather than a direct effect of phenology on within-canopy patterns.

Instead, inter-annual variation remained the more dominant driver of within-canopy caterpillar metrics. Caterpillar densities and diversity were still consistently higher in 2024 than 2023 across canopy levels, with significant positive effects of 2024 on the conditional component for total caterpillar density (β=0.77, 95%CI [0.50, 1.04]), winter moth density (β=0.66, 95%CI [0.34, 0.95]), green oak tortrix density (β=0.90, 95%CI [0.58, 1.22]), and species richness density (β=0.65, 95%CI [0.43, 0.88]). Again similarly to the main level by year models, 2025 was associated with a higher probability of caterpillar absence across most responses, indicated by significant positive effects in the hurdle component for total caterpillar density (β=2.35, 95%CI [0.59, 4.31]), winter moth density (β=1.80, 95%CI [0.63, 3.03]), green oak tortrix density (β=2.80, 95%CI [0.61, 5.62]), species richness density (β=2.17, 95%CI [0.39, 4.03]), and Shannon diversity values (β=1.83, 95%CI [0.65, 3.01]). Herbivory across canopy levels showed no significant relationships with phenology or year. Overall, these results indicate that temporal variation between years exerted a stronger influence on caterpillar communities than variation in host tree phenology, regardless of how phenology was measured.

### Correlation with ground sampling methods

To assess how well ground-based sampling reflected canopy-level caterpillar metrics, we calculated Spearman correlation coefficients between branch-sampled methods and corresponding ground methods (**Table 1**).

**Table 1.** Spearman correlations (r_s_) between ground sampling methods (frass or water traps) and corresponding caterpillar community aspects measured on branch samples, varying across canopy level (2024: lower n=30, mid n=34, upper n=34; 2025: lower n=25, mid n=33, upper n=33). Water trap data here were collected on the closest sampling day to MEWP sampling. Correlation coefficients in bold denote significance at p<0.05; asterisks indicate levels of significance (*=p<0.05, **=p<0.01, ***=p<0.001).

| *Response* | *Level* | *2024* | *2025* | *Response* | *Level* | *2024* | *2025* |
| --- | --- | --- | --- | --- | --- | --- | --- |
| Frass mass ~<br>total<br>caterpillar<br>density | Lower | **0.68\*\*\*** | -0.06 | Frass mass<br>~ herbivory | Lower | **0.37\*** | 0.36 |
|  | Mid | **0.49\*\*\*** | **0.54\*\*** |  | Mid | **0.38\*** | **0.43\*** |
|  | Upper | **0.59\*\*\*** | **0.63\*\*\*** |  | Upper | **0.50\*\*** | **0.36\*** |
| Total<br>caterpillar<br>abundance<br>(water) ~<br>density | Lower | -0.30 | -0.12 | Winter<br>moth<br>abundance<br>(water) ~<br>density | Lower | 0.09 | 0.19 |
|  | Mid | 0.06 | **0.41\*** |  | Mid | 0.31 | **0.36\*** |
|  | Upper | -0.10 | **0.44\*\*** |  | Upper | 0.15 | **0.53\*\*** |
| Green oak<br>tortrix<br>abundance<br>(water) ~<br>density | Lower | -0.24 | NA | Species<br>richness<br>(water) ~<br>density | Lower | -0.18 | -0.25 |
|  | Mid | -0.10 | NA |  | Mid | 0.11 | **0.36\*** |
|  | Upper | -0.23 | NA |  | Upper | 0.04 | **0.42\*** |
| Shannon index<br>(water) ~<br>branches | Lower | **-0.37\*** | -0.31 |  |  |  |  |
|  | Mid | 0.05 | 0.04 |  |  |  |  |
|  | Upper | 0.26 | 0.08 |  |  |  |  |

Frass mass, collected over three days preceding branch sampling, showed consistent and moderately strong positive correlations with total caterpillar density on branches across most canopy levels and years (r_s_=0.49-0.68, p<0.01; **Table 1**). The only non-significant relationship occurred for lower branch densities in 2025 (r_s_=-0.06, p=0.783). This indicates that frass mass is generally a reliable indicator of total caterpillar density within tree crowns, independent of canopy level. Although slightly weaker, frass mass similarly correlated significantly with canopy herbivory (both are measures of community productivity) in the same five groups (r_s_=0.35-0.50, all p<0.05; **Table 1**). As frass is composed of faecal deposition from the entire caterpillar community above, we only analysed its relationship with community level metrics rather than species-specific or diversity aspects.

Relationships between canopy level measures and corresponding water trap data collected on/close to MEWP sampling day were more variable and typically weaker (**Table 1**). In 2024, the only significant correlation occurred between Shannon diversity values on lower branches with water traps, but this relationship was unexpectedly negative (r_s_=-0.37, p<0.05). In 2025, several weak to moderate positive correlations were found: total caterpillar density correlation with total caterpillar abundance in water traps for mid (r_s_=0.41, p<0.05) and upper branches (r_s_=0.44, p<0.01), and similar relationships were detected for winter moth and species richness in the same canopy levels (r_s_=0.36-0.53, all p<0.05). No significant associations were found for Shannon diversity in 2025, and green oak tortrix was excluded from that year due to only one branch-collected individual.

Collectively, these results indicate that frass traps provide a more robust proxy for total caterpillar densities, while water traps capture some elements of canopy-level community aspects, particularly in mid and upper canopy levels, but that may be strongly influenced by sampling year. The more inconsistent relationships for Shannon diversity and green oak tortrix density suggest water traps are less effective for capturing finer-scale community structure and certain taxa (i.e. Shannon accounts for both abundance and species identity of individuals, unlike just species richness).

However, when comparing canopy measures from the single MEWP sampling events against the cumulative total of caterpillars collected in water traps across the whole season, some associations became stronger, particularly for the 2024 estimates (**Table 2**). Total caterpillar abundance was now moderately correlated between methods for 2024 as well as 2025 for mid and upper branches (r_s_=0.37-0.55, all p<0.05). Cumulative winter moth was similarly significantly related to MEWP measures in 2024, with the strength of relationship improving with higher strata, as well as 2025 (r_s_=0.42-0.64, all p<0.05). Upper level species richness in 2024 also became well correlated when using overall seasonal values (r_s_=0.42, p<0.05). However, even when using seasonal totals, correlations remained absent for green oak tortrix density and Shannon diversity, indicating that water traps are still limited in capturing taxa with different life histories and finer-scale community structure.

**Table 2.** Spearman correlations (r_s_) between ground sampling methods (frass or water traps) and corresponding caterpillar community aspects measured on branch samples, varying across canopy level (2024: lower n=30, mid n=34, upper n=34; 2025: lower n=25, mid n=33, upper n=33). Water trap data here were the cumulative totals of caterpillars collected across the whole season. Correlation coefficients in bold denote significance at p<0.05; asterisks indicate levels of significance (*=p<0.05, **=p<0.01, ***=p<0.001).

| *Response* | *Level* | *2024* | *2025* | *Response* | *Level* | *2024* | *2025* |
| --- | --- | --- | --- | --- | --- | --- | --- |
| Total caterpillar abundance (water) ~ density | Lower | 0.18 | -0.11 | Winter moth abundance (water) ~ density | Lower | **0.48\*\*** | 0.11 |
|  | Mid | **0.45\*\*** | **0.53\*** |  | Mid | **0.51\*\*** | **0.42\*** |
|  | Upper | **0.37\*** | **0.55\*\*\*** |  | Upper | **0.64\*\*\*** | **0.53\*\*** |
| Green oak tortrix abundance (water) ~ density | Lower | -0.12 | NA | Species richness (water) ~ density | Lower | 0.22 | 0.03 |
|  | Mid | 0.00 | NA |  | Mid | 0.29 | **0.52\*\*** |
|  | Upper | -0.01 | NA |  | Upper | **0.42\*** | **0.60\*\*\*** |
| Shannon index (water) ~ branches | Lower | -0.23 | -0.19 |  |  |  |  |
|  | Mid | 0.09 | 0.06 |  |  |  |  |
|  | Upper | 0.13 | 0.27 |  |  |  |  |

## Discussion

Across three years, vertical stratification of caterpillar community measures within oak canopies was modest and, notably, inconsistent, with both the direction and magnitude of differences among canopy levels varying between years. Although budburst timing varied systematically within crowns, this fine-scale phenological heterogeneity did not explain variation in caterpillar community measures, indicating that within-canopy differences in leaf emergence have limited influence on later caterpillar distributions. Instead, variation among years and among individual trees had more prevalent effects across community measures although this was not driven by NDVI-derived tree phenology. Of ground-based monitoring approaches, frass mass provided a reliable proxy of canopy caterpillar density and herbivory, whereas water traps showed weaker and more variable relationships with corresponding canopy measures across years.

### Vertical and temporal variation in caterpillar communities

Caterpillar community composition, density and diversity metrics varied weakly to moderately among canopy levels, with some significant differences, but patterns were inconsistent across years (**Figures 1 & 2**). This aligns with other temperate and tropical studies where vertical stratification often varies in strength and direction depending on local contexts (e.g. Seifert et al., 2020; Neves et al., 2013; Brehm, 2007; Blair, 2024; Šigut et al., 2018; Finnie et al., 2024). In our study, total caterpillar, winter moth, and species richness densities increased from lower to upper canopy levels in 2023, reversed in 2024 (when overall densities were higher), and showed no vertical patterns in 2025 when caterpillar densities were generally low (**Figure 2a-b, d**). For winter moths specifically, the increase in densities toward the upper canopy in 2023 is consistent with expectations from their oviposition behaviour and vertical gradients in host phenology (**Figure 4**; Watt et al., 1992), whereas the absence of vertical stratification in general early instar densities in April 2024 (**Figure 3**) indicates that such effects are not consistently expressed around the time of larval emergence (i.e. vertical structure can emerge at peak abundance without being detectable at the earliest larval stages). This may be explained by starvation tolerance (Tikkanen & Julkunen-Tiitto, 2003) or dispersal within or between trees to track optimal food quality, even of early instars, given the suspected preference of winter moth for ovipositing in the upper canopy (Zulin et al., 2018; Carroll & Quiring, 1994; Watt et al., 1992).

**Figure 3.**
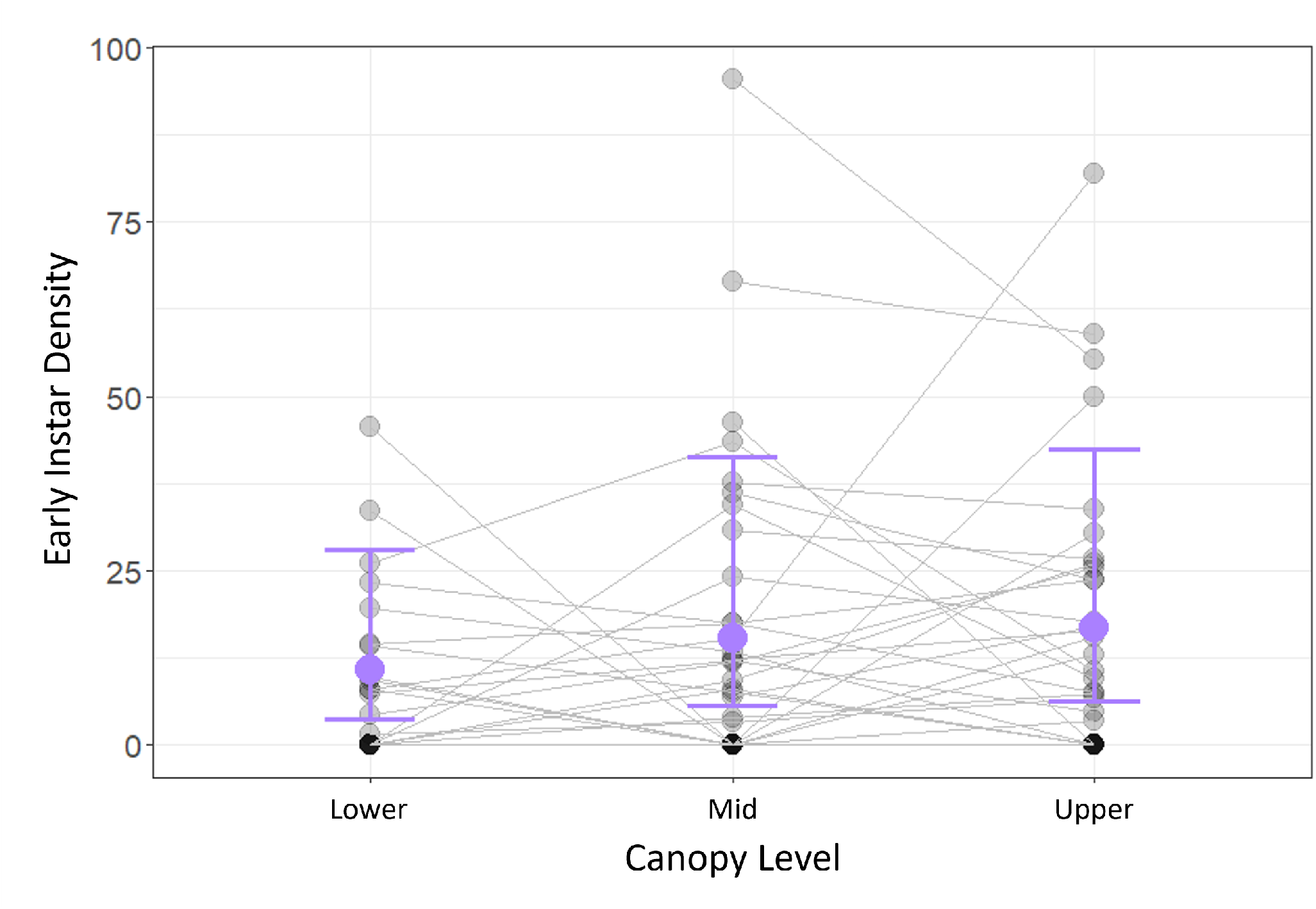
Conditional effects of canopy level (lower, mid, upper third) on early instar caterpillar density across n=92 branches from 34 oak trees in April 2024. Large points and error bars show means and 95% credible intervals from the combined hurdle lognormal model components, averaged across random effects of tree identity (grey lines) and bud stage. Densities are abundance counts scaled to 100g bud mass. No significant (p<0.05) pairwise differences were detected among canopy levels based on model estimated marginal means

**Figure 4.**
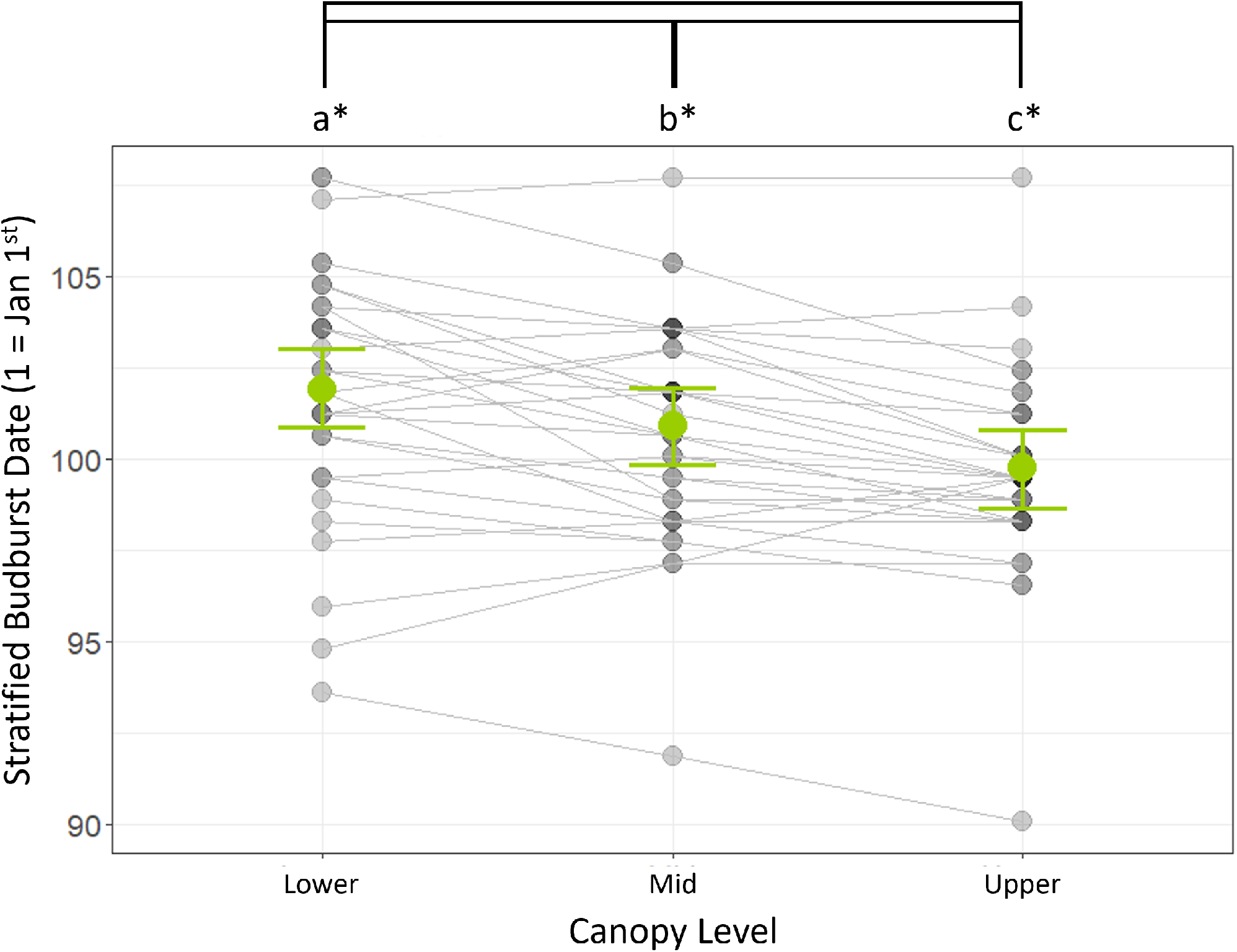
Conditional effect of canopy level (lower, mid, upper third) on stratified budburst dates for n=98 branches from 34 oak trees in 2024. Large green points and error bars show means and 95% credible intervals from a Bayesian linear mixed model, averaged across the random effect of tree identity (grey lines). Letters and asterisks denote significant (p<0.05) pairwise differences between canopy levels based on model estimated marginal means.

Differences among canopy-level means were small relative to total variance, and many branches contained no caterpillars, requiring the hurdle lognormal model structure. These variable modest differences between canopy levels suggest that caterpillar communities are only weakly stratified vertically. This could further indicate some degree of within-tree connectivity, for instance via behavioural dispersal (Carroll & Quiring, 1994; Zulin et al., 2018). Weak to moderate within-level repeatability also implies fine-scale horizontal spatial variability associated with orientation or branch structure. Experimental work has shown differential caterpillar growth between northern and southern canopy aspects, or between proximal versus terminal branch positions, as well as studies documenting densities of invertebrate taxa varying by canopy section (e.g. *Eurytoma* gall wasps, Leite et al., 2009; Suomela & Nilson, 1994; Suomela et al., 1995a; Suomela et al., 1995b). These findings reinforce that canopy complexity may shape caterpillar distributions in multiple dimensions, not just vertical height.

As with density measures, Shannon diversity and herbivory similarly peaked on upper branches in 2023, were higher overall but vertically uniform in 2024, and while Shannon diversity declined in 2025, herbivory remained relatively high that year and was greatest on mid-canopy branches despite reduced caterpillar densities (**Figure 2e-f**). The 2025 results may reflect higher feeding rates per individual, or that sampling occurred after the caterpillar phenological peak, when abundance had begun to decline but herbivory had already accumulated. These inter-annual shifts emphasise the importance of multi-year sampling for detecting dynamic canopy patterns that may or may not persist under annually varying environmental conditions.

### Inter-annual shifts in caterpillar phenology and sampling timing

The contrasting direction of canopy-level density patterns between 2023 and 2024 may be explained by inter-annual differences in the timing of sampling relative to caterpillar phenology (**Figure 2a-b, d**). Our water trap monitoring data show that MEWP sampling in 2023 occurred substantially earlier than the ground-based caterpillar phenological peak (water trap mean = day 147 (n=77 oaks); sampling days = 135-141). At this earlier time point, upper branches – likely to be more phenologically advanced, as shown by approximately two days earlier budburst in the upper canopy in 2024 (**Figure 4**) – may have supported higher caterpillar densities, while mid and lower canopy branches were still approaching peak suitability. In contrast, sampling in 2024 took place much closer to, or during, the estimated phenological peak (water trap mean = day 137 (n=120 oaks); sampling days = 134-139), meaning that caterpillar abundance in the more phenologically advanced upper canopy may already have begun to decline due to pupation or declining leaf quality, while mid and lower branches were reaching peak abundance. The same sampling window in 2025 (days 132-137) occurred slightly later than the ground-based phenological peak (water trap mean = day 132 (n=83 oaks), i.e. caterpillar phenology became earlier each year but trees were sampled during the same calendar week), which could explain both the markedly lower densities remaining on branches and the absence of vertical gradients, as more caterpillars across all canopy levels had completed development. Such inter-annual shifts in phenology relative to fixed calendar sampling are likely to influence apparent vertical caterpillar distributions, something rarely accounted for in studies relying on single-year sampling or limited temporal resolution. This may help explain why vertical stratification patterns in canopy caterpillar communities are often inconsistent across contexts, as patterns may depend on exactly when trees are sampled during the caterpillar phenological distribution.

### Host tree phenology

Host tree phenology is a key driver of caterpillar phenology and abundance and varies both within and between oak canopies, with the potential to influence caterpillar community structure (van Asch & Visser, 2007; Barber & Fahey, 2015). Although we detected significant within-crown heterogeneity in budburst timing in 2024 (**Figure 4**), this small temporal offset appeared to have negligible consequences for caterpillar density, diversity or herbivory. Early instar densities collected in April around the time of budburst showed no vertical stratification (**Figure 3**), and stratified budburst date did not improve model fit relative to whole-crown budburst, suggesting that fine-scale phenological variation within trees does not strongly structure caterpillar communities.

One explanation is that a two day difference in budburst timing, while statistically significant, is short relative to between-tree phenological variation and falls within the starvation tolerance window documented for species such as *O. brumata*, reducing selection for precise synchrony with local leaf emergence (Tikkanen & Julkunen-Tiitto, 2003; Stamp & Bowers, 1990). Early instars may therefore tolerate feeding on sub-optimal buds until budburst progresses locally, or disperse within or between trees to track optimal resources, weakening vertical structuring around larval emergence. Leaf chemical and structural traits are also relatively more similar across sunlit upper and shaded lower canopy positions shortly after budburst, before divergence later in the season (Stamp & Bowers, 1990; Molleman et al., 2022). Consequently, during our mid-May sampling, larvae likely experienced broadly comparable food quality across canopy strata despite the small initial budburst offset, reducing selective pressure for precise fine-scale within-crown phenological synchrony and explaining why measured budburst differences occurring a few weeks earlier did not correspond with vertical patterns in early instar or peak caterpillar community measures.

Additionally, our results also indicate tree identity had a particularly strong effect on budburst phenology (**Figure 4**) i.e. early crowns tend to be early across all canopy levels, further reinforcing that within-crown phenological variation may be significant but small compared to between tree variation, and suggesting that synchrony should mainly operate at the scale of individual oak trees.

However, extending to a whole-crown inter-annual scale using NDVI-derived phenology further indicated that between-tree phenological variation exerted relatively weak effects on caterpillar communities. While half-NDVI dates captured substantial inter-annual advancement of 16 days across study years, caterpillar densities, diversity and herbivory were largely insensitive to variation in host tree phenology once this inter-annual effect was controlled for. There was only limited evidence of species-specific responses for green oak tortrix occurrence. This suggests that, within the range of phenological variation observed, synchrony between caterpillars and host trees may be sufficiently flexible to buffer any mismatches in timing. Although, while correlated with bud-burst date, half-NDVI does not represent the same event and is shifted later than budburst which may explain weaker relationships with caterpillar measures (Morley et al., 2025).

### Alternative or complementary mechanisms

Beyond phenological timing, additional mechanisms may also contribute to vertical canopy gradients in caterpillar communities. For example, feeding guild composition may shift with canopy microclimate, favouring species better adapted to exposed or sheltered conditions, thus influencing community species diversity. For instance, shelter-building tortricids may tolerate the more exposed upper canopy branches by modifying leaves into protective structures (Seifert et al., 2020). However, our analyses focused primarily on taxonomic rather than functional guild diversity (**Figure 1**), and the most abundant leaf-rolling tortricid in our system, *Tortrix viridana*, did not show a strong preference for upper branches (**Figures 1 & 2c**), suggesting that guild-level patterns may vary among taxa and contexts still depending on organisms’ specific niche requirements.

Secondly, increased sunlight in the upper canopy can elevate leaf surface temperatures and potentially accelerate larval development or activity. Sun-exposed leaves may also differ in nutritional and structural traits, including leaf toughness, and carbon, water and tannin content, thereby affecting food quality and caterpillar performance i.e. herbivory levels and survival (Yamasaki & Kikuzawa, 2003; Hakimara & Despland, 2025a; Corff & Marquis, 2001; Hemming & Lindroth, 1995; Eisenring et al., 2021; Blair, 2024; Ruhnke et al., 2009). Stiegel et al. (2017), for example, found low herbivory in upper canopies of beech (*Fagus sylvatica*) due to indirect microclimatic effects of temperature on leaf traits, including higher leaf toughness and carbon content and lower nitrogen availability, as did Hakimara & Despland (2025b) in sugar maples (*Acer saccharum*). This contrasts with our findings of higher herbivory on upper branches in 2023 and 2025, indicating an influence of other mechanisms, but is consistent with the more uniform vertical herbivory observed in 2024 (**Figure 2f**). Conversely, in cooler or wetter years, such as 2024 at our study site, the more sheltered leaves in the lower canopy may become more favourable for caterpillar survival, compounded by rain dislodging caterpillars from exposed leaves. Consistent with this, caterpillar densities and diversity on lower branches were higher in 2024 than on other strata, although densities in mid and upper branches also tended to be greater that year (**Figure 2a-d**), suggesting that favourable early spring conditions may have enhanced larval establishment or synchrony with host phenology at the whole-tree scale.

The very low caterpillar densities observed in 2025, including a near absence of *T. viridana*, likely reflect a combination of factors: population cycles declining towards a trough, reduced survival under drier spring conditions affecting leaf quality or phenological synchrony, in addition to sampling occurring after the caterpillar phenological peak. Together, these observations reinforce evidence from other systems that inter-annual variability in weather, host phenology and food resource quality can outweigh within-tree spatial heterogeneity in structuring caterpillar communities (Corff & Marquis, 2001; Ruhnke et al., 2009).

### Evaluation of ground-based sampling methods

Comparing canopy and ground-based sampling methods highlights biases influencing how well they reflect arboreal caterpillar communities. The moderately strong positive correlations between frass mass and total caterpillar density across canopy levels (r_s_=0.49-0.68, p<0.01 in most cases; **Table 1**) were generally consistent across 2024 and 2025 and canopy strata, confirming that frass traps provide a reliable proxy for overall canopy caterpillar densities, and similarly for their productivity. This agrees with strong correlation between caterpillar biomass estimated from frass traps versus branch sampling in Visser et al. (2006).

Conversely, correlations between water trap abundances and canopy branch data were weaker and less consistent, when using measurements from the closest sampling day, with significant associations limited to mid and upper canopy comparisons in 2025 (**Table 1**). This inconsistency likely reflects methodological biases linked to caterpillar behaviour. Water traps collect larvae that fall from the canopy, either intentionally descending to pupate or accidentally dislodged, so therefore only represent a subset of the canopy community. Species that pupate in the soil, such as winter moths, are consequently well represented and showed significant correlations with mid and upper branch densities in 2025 when sampling dates were aligned (r_s_=0.36, p<0.05, and r_s_=0.53, p<0.01 respectively), with additional seasonal total correlations observed in 2024 (**Table 2**). Meanwhile canopy-pupating or sessile species, such as the green oak tortrix, are underrepresented and showed no significant relationships with branch measures. Hence community-level measures, such as total caterpillar and species richness densities, exhibited only moderate positive relationships with branch measures in 2025 due to a mixture of species life histories (**Tables 1 & 2**). Furthermore, these positive associations between canopy data and water trap measures (on closest sampling day) were only present in 2025, further highlighting the need for multi-year sampling in ecological studies.

However, it was reassuring that the strengths of relationships improved for total caterpillar and winter moth densities, particularly in 2024, when seasonal total abundances were used (**Table 2**). This suggests that single sampling event water trap measures are driven by short-term variability in larval falling rates, meaning that counts on any given day may not accurately reflect the standing density of caterpillars within the canopy. Larval falling and capture rates may be affected by both species-specific behaviours and stochastic environmental factors e.g. high winds and rainfall dislodging individuals. Other caterpillars from non-focal trees could laterally drift into traps, and understorey vegetation could intercept them first (we placed traps under clear areas of canopy as far as possible to minimise this). In contrast, cumulative seasonal totals integrate this noise over time, providing more stable estimates aligning more closely with canopy measures despite being collected over different temporal scales.

Together, this indicates water traps systematically underestimate total arboreal caterpillar densities and diversities on single sampling events, biasing our estimates of canopy community composition and taxa with different life histories. They are still valuable for monitoring phenological timing over the season, if interpreted cautiously, as stochasticity is smoothed over and descending late instars signal the seasonal peak timing even if absolute numbers and species representation are incomplete. Frass sampling offers a stronger quantitative link to total canopy biomass on any given day, regardless of species-specific behaviours, but logistical challenges such as dissolving during rainfall and limited taxonomic resolution remain.

### Limitations and future directions

Although our sampling spanned three years and multiple canopy strata, sampling aimed to occur around the time of peak caterpillar abundance each year. As discussed above, this was not necessarily right on the peak given logistics of hiring machinery. While species-level identifications were possible for dominant taxa like winter moths and green oak tortrix caterpillars, 5.65% of collected individuals remained unidentified to (morpho)species level, potentially underestimating species richness and Shannon diversity. Future work using greater taxonomic resolution, e.g. through metabarcoding, could better capture the full community composition. Quantitative measurements of simultaneous leaf traits (e.g. chlorophyll, moisture or nitrogen content, toughness and thickness) or microclimate would also strengthen our mechanistic understanding of drivers of the variable vertical stratification we observed beyond what we could infer from host tree phenology and early instar densities (e.g. Blair, 2024; Hakimara & Despland, 2025b).

## Conclusions

This multi-year field study shows that vertical stratification in oak canopy caterpillar communities is present but is inconsistent across years, with temporal, environmental, and host tree variation exerting stronger influences on community structure. Inter-annual shifts in caterpillar density, diversity, and herbivory, including reversals in vertical patterns between years, highlight the need for multi-year, phenologically contextualised sampling to disentangle the effects of canopy structure and broader climatic drivers on community dynamics. Modest within-tree vertical variation in budburst appears insufficient to generate vertical stratification in caterpillar communities, suggesting that climate-induced shifts in mean budburst timing will primarily influence caterpillar populations at the between-tree and between-year scales. Consequently, this supports the use of whole-crown phenology metrics, such as drone-derived NDVI curves, for capturing inter-annual trends, while highlighting that such measures alone may have limited predictive power for within-canopy caterpillar distributions. Ground-based frass traps provided a reliable proxy for canopy caterpillar biomass, whereas water traps did not as consistently reflect community patterns due to behavioural and methodological biases. These findings have implications for understanding canopy herbivore dynamics with host trees, designing multi-year ecological studies, and interpreting biases in indirect sampling methods. Integrating canopy-access sampling with robust ground-based proxies such as frass monitoring will enhance predictions of forest resilience and trophic synchrony between host trees and the caterpillar communities they support under on-going climatic or structural change.

## Data Accessibility Statement

The data and analysis scripts that support the findings of this study are available on Zenodo at DOI: https://doi.org/10.5281/zenodo.19453244 (Morley, 2026).

## Author Contributions

LMM, EFC and BCS contributed to the ideas of this study. LMM collected data and conducted the statistical analyses (with inputs on drone data methodology from SJC). SJC led on MEWP sampling in the field. LMM wrote the manuscript with EFC, BCS and SJC providing feedback. All authors approved the final manuscript. LMM: Conceptualization; writing – original draft preparation; investigation; formal analysis; writing – review and editing. EFC: Conceptualization; writing – review and editing. SJC: Resources; software; writing – review and editing. BCS: Conceptualization; writing – review and editing.

## ACKNOWLEDGEMENTS

We thank the many field assistants who helped with data collection, including Léa Beaupère, Gwen Lewis, Amber Winship, Zoe Young, Mia Croft and Sienna Rattigan, and other members of the PhenoScale Group (Department of Biology, University of Oxford) for further assistance in the field and their comments during the development of this work. We also thank the funder (UKRI Frontiers grant EP/X024520/1 to BCS) for supporting this work.

