## Supplementary Information for "Vertical Variation of the Caterpillar Community in Oak (*Quercus robur*) Canopies"

**Contents**

Supplementary Figure 1 (map of sampled trees) ........................................................................................ 2

Supplementary Figure 2 (justification of linearly scaling species richness) ............................................. 3

Supplementary Table 1 (statistical outputs of canopy level and year models on caterpillar community measures) ........................................................................................................................ 4-5

Supplementary Table 2 (statistical outputs of canopy level on early instar density model) .................. 6

Supplementary Table 3 (model comparison statistics of whole-crown versus stratified budburst on caterpillar community measures) ..................................................................................................... 7

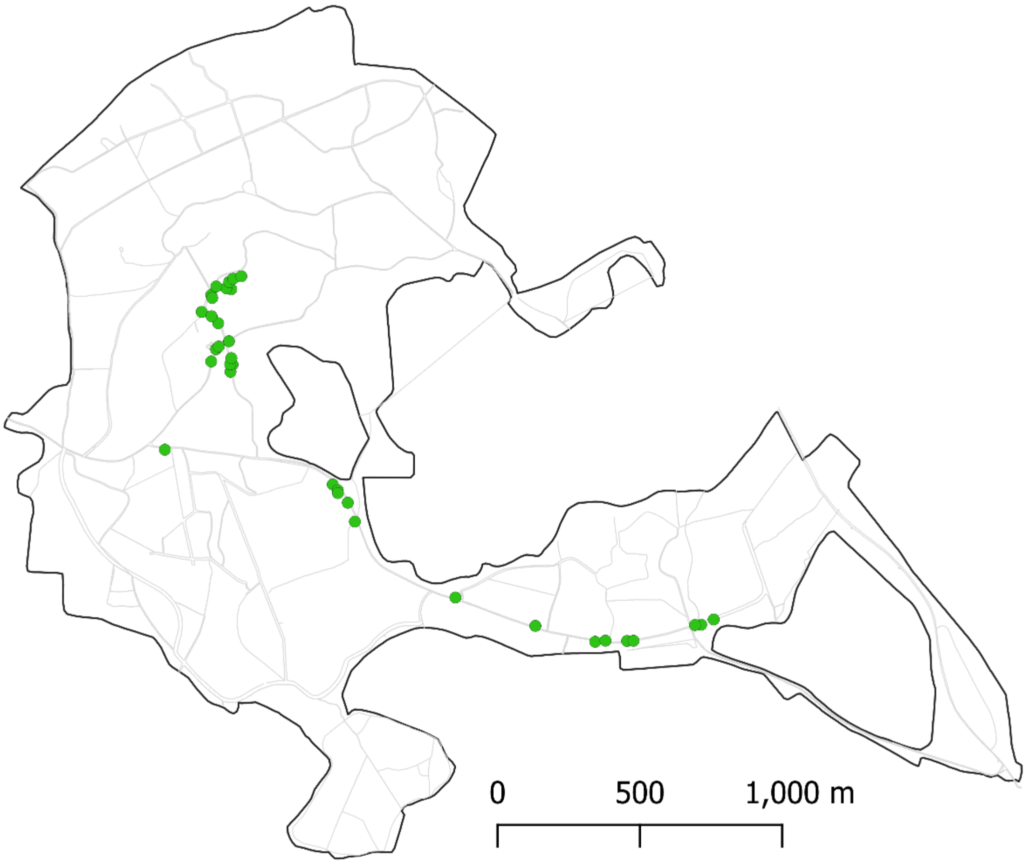


**Supplementary Figure 1.** Map of Wytham Woods showing the locations of 34 oak trees (*Quercus robur*) sampled for within-canopy caterpillar community measures in this study.

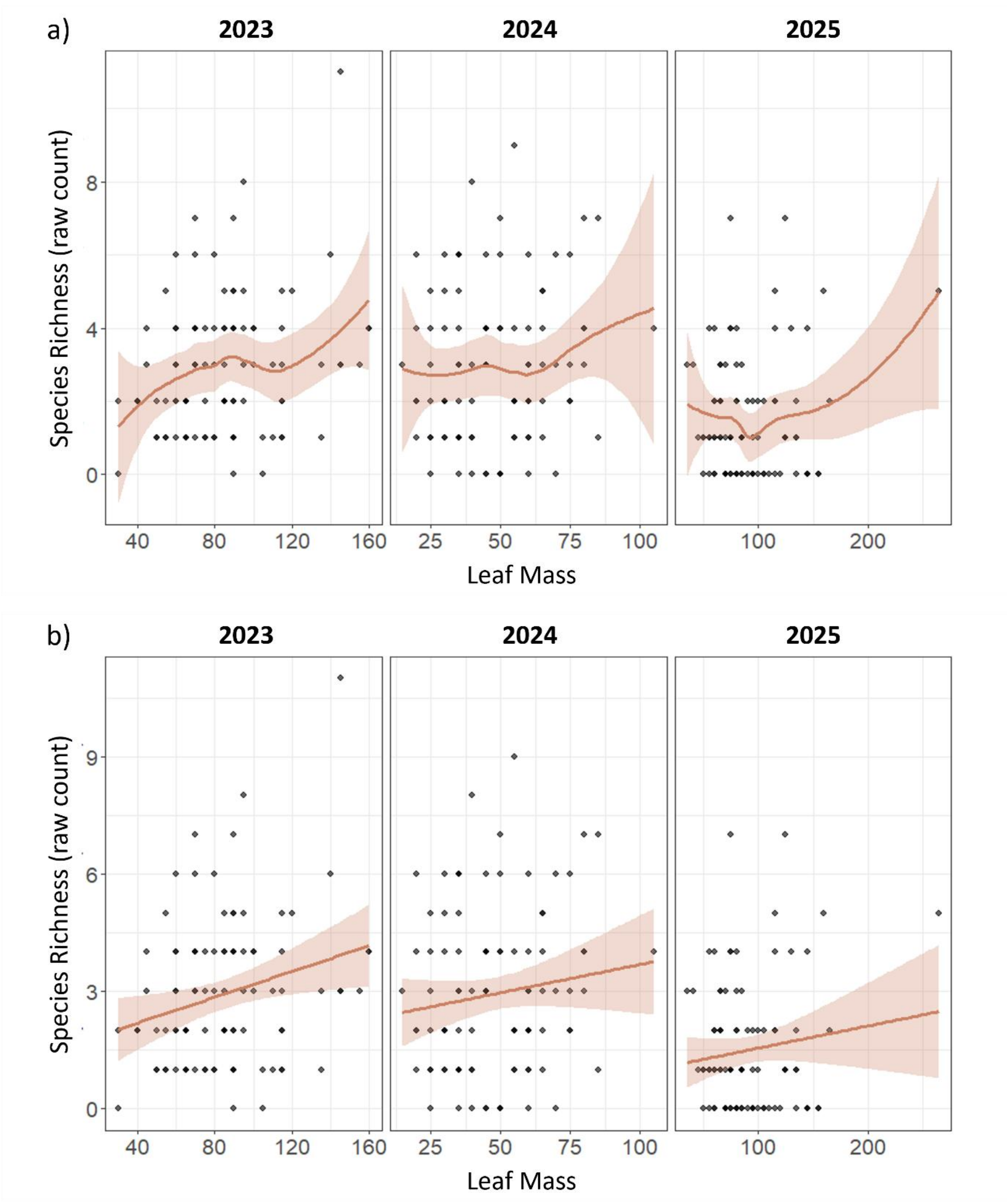


**Supplementary Figure 2.** We evaluated the assumption that species richness scales linearly with sampling effort by plotting raw species richness against branch leaf mass by year: **a)** with a loess smoother and **b)** with a linear regression model. Species richness increased weakly but approximately linearly with leaf mass in all years, with no evidence of saturation across the range of leaf masses sampled, supporting the use of species richness density as counts standardised by leaf mass rather than rarefaction-based approaches.

| | | | | |
|---|---|---|---|---|
| ***Total Caterpillar Density ~ Level x Year*** *(n=285)* | | | | |
| $R^2$ = 43.9% | *Non-zero effects* | *Estimate [95%CI]* | *Zero hurdle effects* | *Estimate [95%CI]* |
| | Intercept | **1.28 [0.95, 1.62]** | Hurdle Intercept | **-2.74 [-4.53, -1.38]** |
| | Mid | 0.28 [-0.11, 0.66] | Mid | -1.16 [-4.33, 1.43] |
| | Upper | **0.61 [0.21, 0.98]** | Upper | **-9.86 [-30.65, -0.42]** |
| | 2024 | **1.14 [0.75, 1.54]** | 2024 | 0.43 [-1.59, 2.52] |
| | 2025 | -0.30 [-0.75, 0.16] | 2025 | **2.15 [0.55, 4.14]** |
| | Mid:2024 | -0.46 [-0.99, 0.08] | Mid:2024 | 1.02 [-2.20, 4.85] |
| | Upper:2024 | **-1.17 [-1.70, -0.64]** | Upper:2024 | 9.22 [-0.70, 30.72] |
| | Mid:2025 | -0.24 [-0.93, 0.28] | Mid:2025 | 0.57 [-2.27, 4.07] |
| | Upper:2025 | -0.54 [-1.14, 0.07] | Upper:2025 | **9.89 [0.37, 31.15]** |
| ***Winter Moth Density ~ Level x Year*** *(n=285)* | | | | |
| $R^2$ = 21.7% | *Non-zero effects* | *Estimate [95%CI]* | *Zero hurdle effects* | *Estimate [95%CI]* |
| | Intercept | **0.74 [0.38, 1.11]** | Hurdle Intercept | -0.58 [-1.37, 0.16] |
| | Mid | 0.11 [-0.31, 0.52] | Mid | **-1.53 [-2.92, -0.23]** |
| | Upper | 0.37 [-0.02, 0.76] | Upper | **-1.91 [-3.50, -0.52]** |
| | 2024 | **0.73 [0.29, 1.15]** | 2024 | -0.47 [-1.60, 0.61] |
| | 2025 | -0.34 [-1.02, 0.37] | 2025 | **2.06 [0.85, 3.40]** |
| | Mid:2024 | 0.14 [-0.44, 0.73] | Mid:2024 | **2.10 [0.44, 3.92]** |
| | Upper:2024 | **-0.75 [-1.31, -0.16]** | Upper:2024 | **2.47 [0.70, 4.39]** |
| | Mid:2025 | 0.10 [-0.73, 0.92] | Mid:2025 | 0.26 [-1.51, 2.09] |
| | Upper:2025 | 0.11 [-0.76, 0.93] | Upper:2025 | 1.29 [-0.66, 3.28] |
| ***Green Oak Tortrix Density ~ Level x Year*** *(n=194)* | | | | |
| $R^2$ = 6.8% | *Non-zero effects* | *Estimate [95%CI]* | *Zero hurdle effects* | *Estimate [95%CI]* |
| | Intercept | **0.72 [0.31, 1.13]** | Hurdle Intercept | 0.60 [-0.18, 1.39] |
| | Mid | -0.21 [-0.73, 0.30] | Mid | -0.35 [-1.41, 0.70] |
| | Upper | -0.01 [-0.49, 0.46] | Upper | **-1.50 [-2.62, -0.44]** |
| | 2024 | **0.98 [0.41, 1.53]** | 2024 | -0.17 [-1.27, 0.87] |
| | 2025 | NA | 2025 | NA |
| | Mid:2024 | -0.22 [-0.97, 0.53] | Mid:2024 | 0.69 [-0.77, 2.18] |
| | Upper:2024 | -0.30 [-0.99, 0.36] | Upper:2024 | 1.31 [-0.11, 2.84] |
| | Mid:2025 | NA | Mid:2025 | NA |
| | Upper:2025 | NA | Upper:2025 | NA |
| ***Species Richness Density ~ Level x Year*** *(n=285)* | | | | |
| $R^2$ = 41.5% | *Non-zero effects* | *Estimate [95%CI]* | *Zero hurdle effects* | *Estimate [95%CI]* |
| | Intercept | **1.02 [0.77, 1.28]** | Hurdle Intercept | **-2.79 [-4.68, -1.43]** |
| | Mid | 0.09 [-0.23, 0.39] | Mid | -1.08 [-4.45, 1.54] |
| | Upper | 0.27 [-0.03, 0.58] | Upper | **-9.97 [-31.48, -0.29]** |
| | 2024 | **1.05 [0.72, 1.37]** | 2024 | 0.85 [-1.01, 3.03] |
| | 2025 | -0.27 [-0.64, 0.11] | 2025 | **2.21 [0.54, 4.28]** |
| | Mid:2024 | -0.30 [-0.74, 0.15] | Mid:2024 | 0.56 [-2.68, 4.38] |
| | Upper:2024 | **-0.84 [-1.27, -0.39]** | Upper:2024 | 9.46 [-0.57, 30.96] |
| | Mid:2025 | -0.07 [-0.57, 0.42] | Mid:2025 | 0.66 [-2.32, 4.21] |
| | Upper:2025 | -0.18 [-0.67, 0.32] | Upper:2025 | **10.12 [0.27, 31.50]** |
| ***Shannon Diversity Index ~ Level x Year*** *(n=285)* | | | | |
| $R^2$ = 18.1% | *Non-zero effects* | *Estimate [95%CI]* | *Zero hurdle effects* | *Estimate [95%CI]* |
| | Intercept | **-0.19 [-0.36, -0.02]** | Hurdle Intercept | -0.60 [-1.39, 0.13] |
| | Mid | 0.16 [-0.06, 0.37] | Mid | -0.63 [-1.75, 0.43] |
| | Upper | **0.34 [0.13, 0.56]** | Upper | -1.02 [-2.30, 0.17] |
| | 2024 | **0.23 [0.01, 0.46]** | 2024 | -0.46 [-1.56, 0.69] |
| | 2025 | 0.05 [-0.25, 0.35] | 2025 | **1.38 [0.26, 2.59]** |

| | | | | |
|---|---|---|---|---|
| | Mid:2024 | -0.15 [-0.46, 0.15] | Mid:2024 | 0.64 [-0.92, 2.23] |
| | Upper:2024 | **-0.35 [-0.66, -0.05]** | Upper:2024 | 1.31 [-0.25, 2.94] |
| | Mid:2025 | -0.16 [-0.52, 0.22] | Mid:2025 | 0.15 [-1.46, 1.69] |
| | Upper:2025 | -0.20 [-0.58, 0.19] | Upper:2025 | 0.68 [-0.96, 2.33] |

| ***Herbivory ~ Level x Year*** *(n=285)* | | |
|---|---|---|
| $R^2$ = 34.8% | *Effects* | *Estimate [95%CI]* |
| | Intercept | **0.53 [0.22, 0.84]** |
| | Mid | 0.13 [-0.26, 0.51] |
| | Upper | **0.73 [0.35, 1.11]** |
| | 2024 | **0.54 [0.15, 0.93]** |
| | 2025 | **0.44 [0.02, 0.85]** |
| | Mid:2024 | 0.03 [-0.51, 0.56] |
| | Upper:2024 | **-0.77 [-1.31, -0.22]** |
| | Mid:2025 | 0.29 [-0.25, 0.85] |
| | Upper:2025 | -0.48 [-1.04, 0.07] |

**Supplementary Table 1.** Outputs of Bayesian mixed models for the effects of canopy level (lower, mid, upper third), year, and their interactions on caterpillar community metrics across n=285 branches from 34 oak trees, sampled from Wytham Woods, UK, in 2023-2025. All density metrics and Shannon diversity index were modelled as hurdle lognormal models to account for zero-inflation (non-zero effects are conditional on caterpillar presence, whilst zero hurdle effects represent the probability of caterpillar absence). Herbivory, which contained no zero values, was modelled using a lognormal distribution. All models included the random effect of tree identity to account for repeated sampling of branches within trees and across years. Intercept was taken as level = lower and year = 2023. 95%CIs are 95% credible intervals around the estimate and effects in bold are significant where CIs do not overlap zero. $R^2$ is the amount of variation explained by the model, including both fixed and random effects. For green oak tortrix density, 2025 was excluded from modelling due to only one individual green oak tortrix larva being recorded which prevented convergence if included in the model. Coefficients are presented on the output link scale, i.e. not back-transformed, for simpler interpretation (i.e. positive coefficients are increasing effects, negative coefficients are decreasing effects, rather than back-transformed to multiplicative effects).

***Early Instar Density ~ Level*** *(n=92)*

| $R^2$ = 36.2% | *Non-zero effects* | *Estimate [95%CI]* | *Zero hurdle effects* | *Estimate [95%CI]* |
|---|---|---|---|---|
| | Intercept | **2.73 [1.75, 3.68]** | Hurdle Intercept | -0.35 [-1.18, 0.46] |
| | Mid | 0.27 [-0.18, 0.73] | Mid | -0.26 [-1.35, 0.83] |
| | Upper | 0.30 [-0.14, 0.78] | Upper | -0.40 [-1.49, 0.69] |

**Supplementary Table 2.** Outputs of a Bayesian mixed model for the effect of canopy level (lower, mid, upper third) on early instar caterpillar density across n=92 branches from 34 oak trees in April 2024. Early instar density was modelled as a hurdle lognormal model to account for zero-inflation (non-zero effects are conditional on caterpillar presence, whilst zero hurdle effects represent the probability of caterpillar absence). The model included the random effects of tree identity and bud stage of the sampled branch. Intercept was taken as level = lower. 95%CIs are 95% credible intervals around the estimate and effects in bold are significant where CIs do not overlap zero. $R^2$ is the amount of variation explained by the model, including both fixed and random effects. Coefficients are presented on the output link scale, i.e. not back-transformed, for simpler interpretation (i.e. positive coefficients are increasing effects, negative coefficients are decreasing effects, rather than back-transformed to multiplicative effects).

| *Response* | *Budburst Type* | *$R^2$ (%)* | *LOOIC [SE]* | *ΔLOOIC (vs best)* |
|---|---|---|---|---|
| Total Caterpillar Density | Whole-Crown | 65.1 | 633.5 [19.9] | +0.6 |
| | Stratified | 64.9 | 632.9 [20.7] | 0.0 |
| Winter Moth Density | Whole-Crown | 43.9 | 453.6 [23.8] | 0.0 |
| | Stratified | 47.1 | 455.1 [24.7] | +1.5 |
| Green Oak Tortrix Density | Whole-Crown | 13.3 | 335.1 [33.2] | +4.5 |
| | Stratified | 17.4 | 330.6 [32.7] | 0.0 |
| Species Richness Density | Whole-Crown | 46.4 | 547.5 [19.3] | 0.0 |
| | Stratified | 45.4 | 548.1 [19.2] | +0.6 |
| Shannon Diversity Index | Whole-Crown | 13.0 | 195.7 [12.1] | 0.0 |
| | Stratified | 13.9 | 196.3 [12.3] | +0.6 |
| Herbivory | Whole-Crown | 64.2 | 415.2 [20.4] | +0.7 |
| | Stratified | 65.4 | 414.5 [20.5] | 0.0 |

**Supplementary Table 3.** Comparison of conditional $R^2$ (proportion of variance explained) and LOOIC values for Bayesian mixed models testing the effect of host tree phenology on caterpillar community metrics across n=98 branches from 34 oak trees in 2024, using budburst date estimated either for the whole-crown or stratified by canopy level (lower, mid, upper third). As before, hurdle lognormal models were used for density and Shannon diversity index responses, and lognormal models for herbivory, with budburst date, canopy level, and their interactions as fixed effects and tree identity included as a random effect. ΔLOOIC values were calculated relative to the best-supported model for each response variable (set to 0), with small ΔLOOIC values (<5) relative to their standard errors indicating no meaningful differences in model fit.
